# Functional Diversification of Tripeptidyl Peptidase and Endopeptidase Sedolisins in Fungi

**DOI:** 10.1101/167379

**Authors:** Facundo Orts, Arjen ten Have

## Abstract

Sedolisins are acid proteases that are related to the basic subtilisins. They have been identified in all three superkingdoms but are not ubiquitous, although fungi that secrete acids as part of their lifestyle can have up to six paralogs. Both tripeptidyl peptidase (TPP) and endopeptidase activity have been identified and it has been suggested that these correspond to separate subfamilies.

We studied eukaryotic sedolisins by computational analysis. A maximum likelihood tree shows three major clades of which two contain only fungal sequences. One fungal clade contains all known TPPs whereas the other contains the endosedolisins. We identified four cluster specific inserts (CSIs) in endosedolisins, of which CSIs 1, 3 and 4 appear as solvent exposed according to structure modeling. Part of CSI2 is exposed but a short stretch forms a novel and partially buried α-helix that induces a conformational change near the binding pocket. We also identified a total of 12 specificity determining positions (SDPs) divided over three SDP sub-networks. The major SDP network contains eight directly connected SDPs and modeling of virtual mutants suggests a key role for the W307A or F307A substitution. This substitution is accompanied by a group of four SDPs that physically interact at the interface of the catalytic domain and the enzyme’s prosegment. Modeling of virtual mutants suggests these SDPs are indeed required to compensate the conformational change induced by CSI2 and the A307. The additional major network SDPs as well as the two small SDP networks appear to be linked to this major substitution, all together explaining the hypothesized functional diversification of fungal sedolisins.

**Highlights:**

There are two sedolisin subfamilies in fungi: tripeptidyl peptidases and endopeptidases
Functional Diversification of fungal sedolisins led to a conformational change in the pocket
Functional Diversification centers around buried SDP307
SDP307 is aromatic in TPPs and Alanine in endosedolisins
Additional SDPs affect the interaction between core and chaperone-like prosegment

## 1 Introduction

### 1.1 Proteases are ubiquitous enzymes that can be classified in many ways

Proteases or peptidases degrade proteins by hydrolysis of peptide bonds. They are involved in various biological processes as cell death [1], nutrition [2] and infections [3]. Proteases are classified, primarily based on the catalytic mechanism, into several classes among which aspartic, cystein, metallo and serine proteases. These main classes are further hierarchically organized into clans and families based on homology [4]. Proteases can also be classified based on other characteristics. A major difference can be made between endo– and exopeptidases, where the latter include di– and tripeptidyl peptidases. Among exopeptidases one can discriminate carboxy– and aminopeptidases.

### 1.2 Sedolisins are acid proteases related with the basic subtilisins

The most common proteases are serine proteases in which a serine serves as the nucleophylic amino acid in the catalytic site. The catalytic site is most often formed by a triade which can differ among the different superfamilies. Currently 12 clans or superfamilies with 29 families have been assigned by MEROPS [5]. One of the most important clans is the SB clan that contains the ubiquitous subtilisins, which include the kexins (S8) [6], and the rather rare sedolisins (S53, for review see [7]). Where the basic subtilisins have a triade that consists of, besides the Serine, a Histidine and an Aspartate, the acid sedolisins have a Glutamate replacing the the Histidine [8]. Despite the different triade and consequently a large difference in optimal activity pH, there is ample evidence the two subfamilies form a superfamily. In 2001 the first structure of a sedolisin, endosedolisin PSCP from *Pseudomonas* sp. 101, complexed with inhibitor iodotyrostatin, was resolved (PDB code 1GA4)

[9], shortly followed by a structure from kumamolysin from Bacillus sp. MN-32 (PDB codes 1T1E for precursor and 1GT9 for mature peptidase [10]). Although sequence similarity between sedolisins and subtilisins is low, structural alignments clearly indicate they are homologous [11] since they have similar folds. Catalytic residues as well as the oxyanion Aspartate appear as homologous. Sedolisins also have a Calcium binding site albeit at a different position than subtilisins [11].

The best studied sedolisin is Human lysosomal CLN2 since mutant forms are involved in the fatal classical late-infantile neuronal ceroid lipofuscinosis or Batten disease. The structure (PDB codes 3EDY for precursor and 3EE6 for mature peptidase) of this tripeptidyl aminopeptidase has been determined and a number of publications describe the effect of many mutations found [12] [13,14]. Of particular interest is W542 which has been shown to be required for activity. The W542L mutant was shown to be retained in the ER which suggests misfolding [15]. In addition, W290L and W307L showed largely reduced activities.

### 1.3 Sedolisins have a large prosegment that appears to have various functions

Also the processing of subtilisins and sedolisins is similar. Both have a large and similar prosegment that appears to be able to form an independent domain that seems to be involved in correct folding of the core or the catalytic domain [16] [17]. The prosegment and catalytic domain are separated by a short propeptide or linker that is removed during isozyme activation at low pH. In general it has been shown that prosegments act in assisting in refolding as well as targeting (for review see [18]). For Human TPP it has been shown that prosegment and catalytic domain have multiple molecular interactions including salt bridges and hydrogen bonds, covering 15% of the solvent accessible surface of the catalytic domain [19]. It has been shown that the prosegment of Human TPP also functions as inhibitor [16]. Secretome analysis of for instance B. cinerea has shown certain paralogs consist of the core part of the enzyme only [20].□

### 1.4 Fungal Sedolisins can have endo- or tripeptidyl-peptidase activity

Other characterized eukaryotic sedolisins are of fungal origin. Scytalidolisin [21], Grifolisin [22] and Aorsin were the first fungal enzymes characterized as sedolisin. More recently, four paralogs from *Aspergillus fumigatus* were characterized. SED_A was, similarly to Aorsin form *Aspergillus oryzae*, characterized as an endosedolisin, whereas SED_B, SED_C and SED_D were shown to have TPP activity [23]. The authors suggested furthermore endosedolisins cluster in a different clade than TPPs, suggesting that gene duplication has resulted in functional diversification. A recent genome paper of fungal plant pathogens *Botrytis cinerea* and *Sclerotinia sclerotiorum* showed that acid secreting fungi as phytopathogens *B. cinerea*, *S. sclerotiorum* but also the saprophytic *Aspergilli* show relatively few subtilisins and many sedolisins, as compared to non-acid secreting fungi such as *Giberella zeae* [24]. This also suggests functional diversification has occurred. Here we study the functional redundancy and diversification of fungal sedolisins by computational analysis. We reconstructed a phylogenetic tree that, together with the underlying multiple sequence alignment (MSA), was used for the identification of cluster specific inserts (CSIs) and specificity determining positions (SDPs), which form the sequence characteristics that can explain functional diversifications. Modeling of wild type and mutant sequences was performed in order to show how functional diversification into endosedolisins and TPP has likely occurred, demonstrating important roles for part of CSI2 and the position homologous to Human TPPs W307.

## 2 Materials y methods

### 2.1 Sedolisin sequence identification

The MEROPS [5] holozyme sequences from the MEROPS database were used to build a first HMMER [25] profile that was used to search a total of 242 complete proteomes as well as the PDB and SwissProt databases. The obtained sequences were scrutinized using MEROPS Batch BLAST [26] which includes a scrutiny for the presence of catalytic site and oxyanion residues. Confirmed sedolisin sequences were aligned and the resulting MSA was used for a second HMMER screening. This process was iterated until convergence.

### 2.2 MSA and Phylogeny

MSAs were constructed using MAFFT’s [27] iterative refinement method and represented using Endscript [28] [29]. The final alignment was manually corrected using as criteria that secondary structure elements (taking as reference 3EE6) should be represented by each sequence combined with entropy minimization. The MSA was trimmed using BMGE [30] (-g 0.3 and h 0.8) which resulted in a trimmed MSA that maintains more than 90% of the columns corresponding to the β-sheets. Maximum likelihood phylogeny was constructed using PHYML3.1 [31] using the WAG model of amino acid substitution using a discrete gamma model of 4 categories and a shape parameter of 1.4, as determined by prior ProtTest [32] analysis. Statistical support was determined by PHYML’s aLRT as well as boostrapping (n = 1024) using the resources available at https://www.e-biothon.fr. The major graphical representations were made using Dendroscope [33]. The presentations showing the CSIs were made using iTOL [34].

### 2.3 Identification of Specifity Determining Positions (SDPs)

We identified CDPs using SDPfox [35]. MISTIC [36] was used to determine levels of mutual information (MI) between positions or columns of the MSA. Initially, CDPs are accepted as SDP when they contain at least two direct connections with other CDPs, using MISTIC’s default z-score cut-off of 6.5. CDPs with a single direct connection are considered as putative SDPs (pSDP) and require additional evidence in order to become accepted as SDP. Cytoscape [37] was used to identify and draw networks of directly connected SDPs. Sequence Logo’s were made using Weblogo [38].

### 2.4 Structure analysis

3D structures of sedolisins were obtained from the Protein Data Bank [39]. 3EE6 [19], corresponding to mature Human TPP was used as a standard. Models were made using I-Tasser [40] using either default settings or using 3EE6 as the reference model. Binding pockets were predicted using Coach [41]. Visualization was performed using VMD [42] which included structural alignment using the STAMP [43] extension. The pocket predictions for 3EE6 and the SED_A and SED_B models were performed with the software Fpocket [44] using the default parameters.

## 3 Results

### 3.1 Datamining, Multiple Sequence Alignment and Phylogeny

In order to perform structure-function prediction of eukaryote sedolisins we set out to obtain a representative collection of sequences. A database containing the complete proteomes of 56 fungi and 186 non fungal eukaryotes (Listed in S. Datafile 2) complemented with the PDB and Swissprot database was subjected to iterative HMMER profiling yielding 230 sequences of sedolisin homologs. Upon hallmark and structural scrutiny a total of 204 sequences were considered as high fidelity, as required for the identification of specificity determining positions. Sequences were aligned and an excerpt of the final MSA is shown in Fig. 1. In general, eukaryotic sedolisins are largely conserved, including the prosegment part. Interestingly, of the three disulfide bridges identified in the resolved structure 3EE6, only the second appears to be conserved among eukaryotes. A trimmed MSA, lacking low quality subalignments, was used to reconstruct a maximum likelihood tree.

**Fig. 1.**
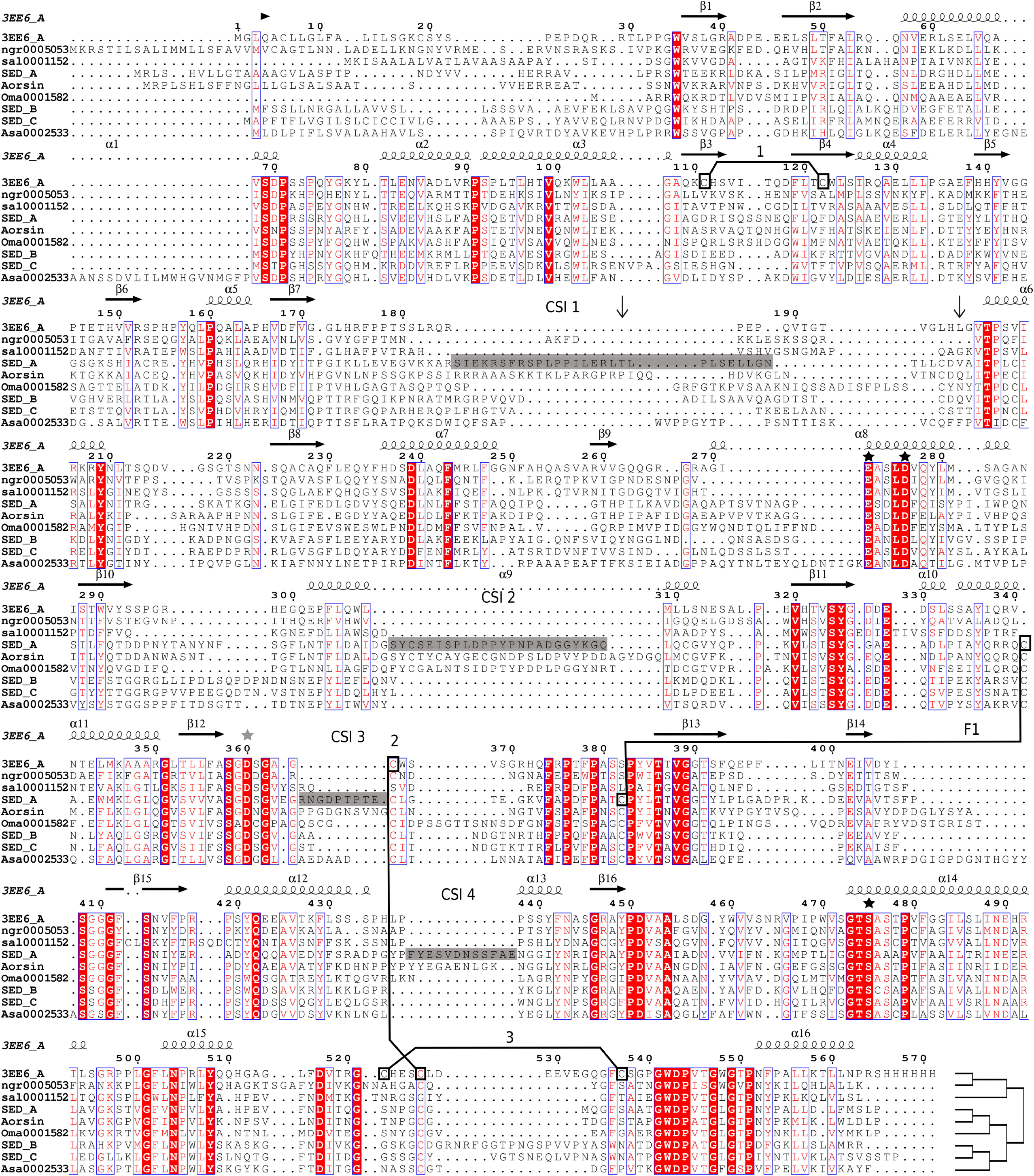
Excerpt of Eukaryotic Sedolisin Sequence Alignment. The demonstrated sequences are representatives of the three mayor phylogenetic clusters, as indicated by the tree placed at the end of the alignment, extracted from the complete MSA (S. Datafile 1). 3EE6, ngr5053 and sal1152 are non-fungal sequences; SED_A and Aorsin are fungal endosedolisins; SED_B and SED_C are fungal TPPs. Oma1582 and Asa2533 are additional fungal sequences with unknown characteristics. Horizontal arrows and helices indicate sheet and helix regions, respectively. Nomenclature and numbers according to 3EE6. The vertical arrows represent beginning and end of the peptide linker. Sequence in gray box represents the SED_A inserts (numbers according to SED_A sequence): CSI1 (190-217), CSI2 (345-370), CSI3 (427-435) and CSI4 (513-524). Black stars indicate catalytic residues E, D, and S whereas the gray star indicates the oxyanion D. Connected boxes, numbered I to III, indicate disulfide bridges identified in the 3EE6 structure. F1 points to two cysteines strictly conserved among all fungal sedolisins but absent in non-fungal sedolisins.

The tree (Fig. 2A) shows three major clades of which two contain only fungal sequences and one contains only non-fungal sequences. Inside the non-fungal clade, vertebrate sedolisins appear in a single subclade that on its turn consists of a mammalian and Aptinopterygii subclade. The mammalian subclade contains human TPP. The apparent random taxonomic distribution of the fungal sequences over the two well separated clades indicates the bifurcation is caused by a functional diversification. In correspondence with Reichard and coworkers [23], one of the fungal clades contains SED_A from A. *fumigatus* and Aorsin [45] from A. oryzae that have both been characterized as endopeptidases. The other fungal clade contains SED_B, SED_C and SED_D from A. *fumigatus*, all characterized as TPPs [23]. Although biochemical evidence is scarce, the above mentioned data suggest sequence diversification has resulted into endo- and TPP sedolisins, hence for the remainder of the manuscript we will refer to the sequences and clades as Hypo-endosedolisin and Hypo-TPP. Both endo and TPP activity has been shown for non-fungal sedolisins, hence, as such we have no indication regarding the state of the ancestral enzyme. In addition, at least Human TPP has been shown to have endo activity at low pH [46].

### 3.2 Cluster Specific Inserts

The MSA demonstrates the presence of four Cluster Specific Inserts (CSIs) we identified in the Hypo-endosedolisins. Fig. 2B shows the distribution of insert length on the phylogeny thereby demonstrating an intricate clustering pattern. CSI2 has about 25 amino acids and is present in all Hypo-endosedolisins whereas CSI4 has about 40 amino acids and is found only in the large subclade of the Hypo-endosedolisin clade. Both show moderate levels of conservation (Fig. 2C). CSI3 is present in all Hypo-endosedolisins as well as in certain Hypo-TPPs and some non-fungal sedolisins. CSI1 is present in all sedolisins but is longer in the Hypo-endosedolisins. CSIs 1 and 3 show no clear conservation.

Because structure determines function we wondered if the CSIs would interfere in the core structure of the protein or if they might appear as solvent exposed loops. We created a model of SED_A and structurally aligned it with 3EE6 (Fig. 3.). Overall, the model is very similar to the 3EE6 structure and the non-homologous CSIs 1, 3 and 4 are indeed predicted at the surface of the mature protein. CSI2 is largely exposed but forms a partially buried helix as is discussed below. This confirms that the CSIs not necessarily affect the basic functional fold of subtilisin-like proteases. Although in the absence of homologous template the prediction of the loop structures formed by the CSIs is not very reliable, the model does allow some general predictions. CSI1 is located next or into the prosegment and likely does not form part of the mature enzyme but rather forms part of the propeptide or linker region (See Fig. 1.). CSI2 and CSI4 are located on opposite ends of the predicted binding pocket (Not shown). CSI3 appears near the calcium binding site.

**Fig. 3.**
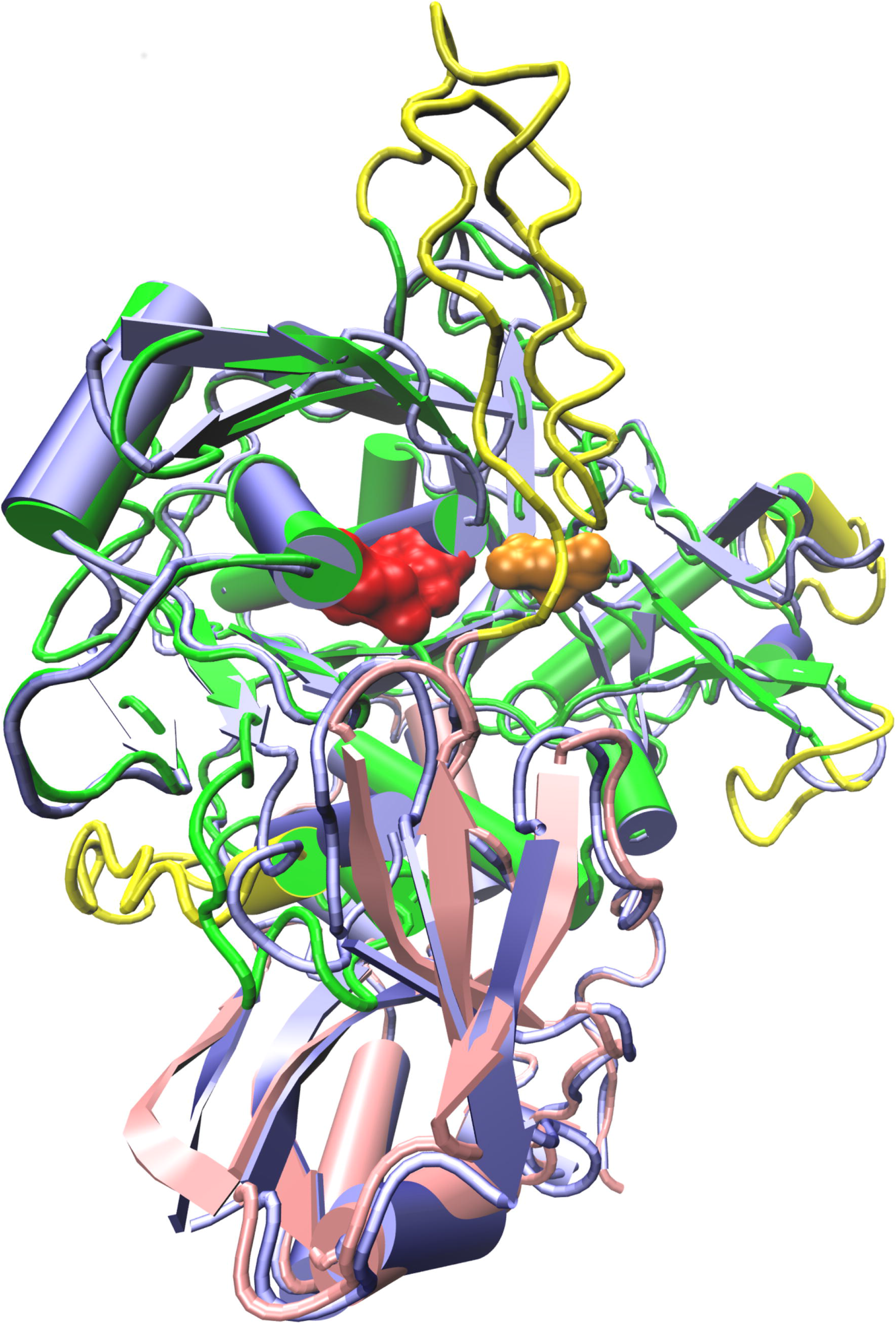
The Cluster Specific Inserts are Solvent exposed. Structural alignment of the 3EE6 structure, represented in blue cartoon and the SED_A model represented in pink (prosegment) and green cartoon (core) with the CSI1 to 4 in yellow cartoon. The red and the orange spheres represent the catalytic residues and oxyanion respectively.

Fig. 4. shows the conformational differences between 3EE6 and the wild type (WT) models SED_A and SED_B as well as a number of mutations that are discussed in the SDP section further below. An important conformational difference is the additional α-helix found in the SED_A model, referred to as H9b, that partially originates from CSI2 (See Fig. 4E). As a result the location of Helix 9 also slightly differs. This seems to have an effect on the binding pocket, as can be seen by a comparison of predicted pockets of SED_B (Fig. 4D) and SED_A (Fig. 4E). The virtual mutant lacking CSI2 does not show helix 9b and retained a structure similar to 3EE6.

**Fig. 4.**
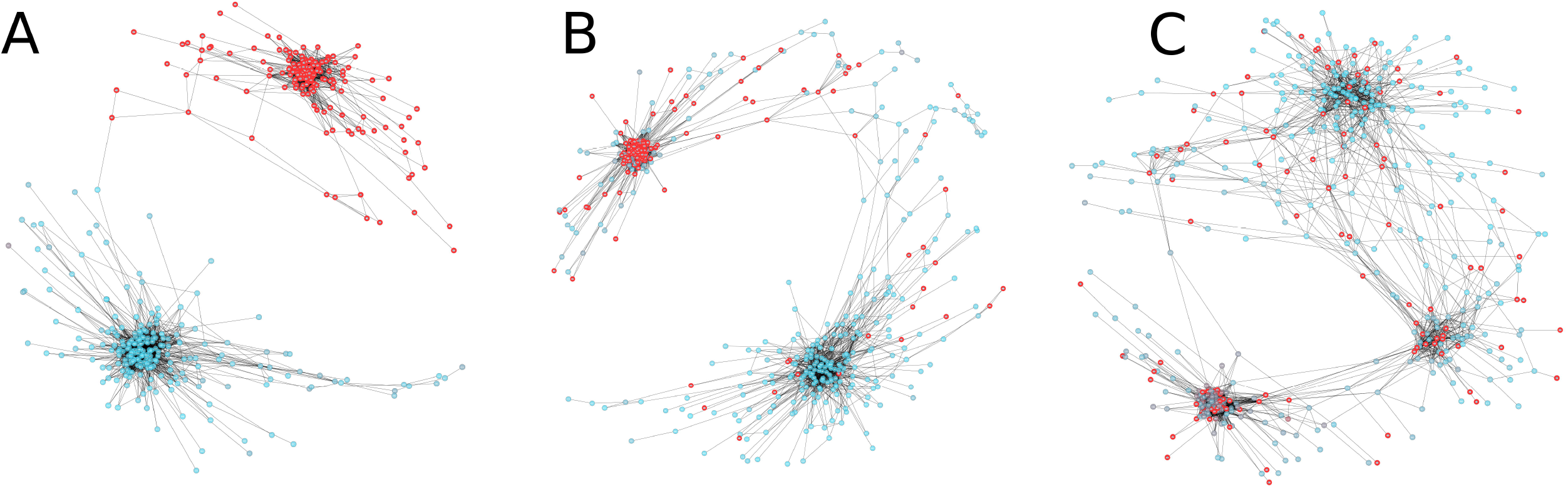
Conformational Differences among Wild Type and Mutant Sedolisins. (A) Cartoon of the SED_A model with regions that have a resQ score below 5 Å in grey homologous regions above 5 Å in yellow and CSIs with resQ above 5 Å in red. (B) ResQ plot of SED_A Modeling (C) Schematic representation of the major structural differences in secondary structure elements of the polypeptide between helices 9 and 12 observed in WT and mutant models of SED_A and SED_B as compared to the 3EE6 structure. Quadruple refers to the virtual SED_A A343F-F92L-K407Q-L410S mutant. Yellow diamonds indicate conserved cysteines that are possibly involved in a novel disulfide bridge. Approximate positions of SDPs 307, 340, 343 and 349 are indicated from left to right by a green check mark. Absent counterparts are represented as a red cross. Helices and sheet are in red cylinders and blue arrows. (D) Local detail of the structural alignment of 3EE6 (blue cartoon) with model obtained for WT SED_B (cyan cartoon). (E) Local detail of the structural alignment of WT SED_A model (yellow cartoon) and the SED_A quadruple mutant model (green cartoon). Helix numbers are indicated in the 3EE6 structure.

### 3.3 SDP identification

We used SDPfox [35] to identify CDPs, positions that contribute significantly to the underlying clustering. We identified 25 CDPs between the Hypo-endo and the Hypo-TPP clusters. Then we performed an analysis of mutual information between positions using MISTIC [36]. Mutual information expresses levels of covariation and high levels suggest co-evolution. CDPs might result form genetic drift but CDPs that show high levels of interaction are more likely specificity determining positions (SDPs). We envisaged that the functional characteristics of phylogenetically well separated subfamilies, such as the Hypo-endo and Hypo-TPP sedolisins, are the result of the interaction of multiple positions. Since therefore, it is likely that one or possibly more subnetworks of directly connected CDPs exist, we accepted all CDPs that connected directly to at least two other CDPs with a score higher than MISTIC’s threshold of 6.5 as SDP. CDPs with a single connection were initially considered as pSDP. Such eventual subnetworks of directly connected CDPs are considered Specificity Determining Networks (SDNs) that not only substantiate that CDPs are SDPs but also show which positions have co-evolved towards a certain diversification.

Next we considered how evolution might have occurred. As stated before, we have no insight in the exact functional characteristics of the ancestral protein. This might have been an endo-sedolisin, a TPP or a protease capable of both reactions. On the one hand we must assume the diversification process in one clade has been largely independent from that in another clade. On the other hand we can also envisage that the diversification processes, although strictly independent, might affect the same positions. As a result, *a priori* one does not know which dataset should be used to determine MI levels. We first performed a global MI analysis comparing the networks obtained when using only the Hypo-endo, the Hypo-TPP and all fungal sequences combined, respectively. Fig. 5. shows that when separate clades are used as dataset, similar networks with partitions in two major clusters are identified. When all fungal sequences are used the MI network partitions into three major clusters, showing little similarity to the partitions of the separate subfamily MI analyses. Hence, it is more likely that the diversification has been the result of independent processes and that as such MI analysis for the identification of SDNs involved in the fungal sedolisin diversification should be determined using clade specific datasets. Table 1 shows the result of the analysis and shows the numbering of SDPs and pSDPs (according to reference sequence of 3EE6) in SED_A and SED_B. We identified an SDN with eight directly connected CDPs using the MI analysis of the Hypo-endo sequences (Table 1, Fig. 6.). The high level of connection suggests these form an SDN of co-evolving SDPs that would correspond with the subfamily´s functional diversification towards and endopeptidase. Only CDP260 was initially considered a pSDP. In addition we identified a pair of connected CDPs that we also initially considered as pSDPs. The Hypo-TPP MI analysis identified a three node SDN that seems unrelated to the Hypo-endosedolisin SDPs, confirming that the diversification processes in both subfamilies are independent and that in this case, as such, for the determination of SDNs it is necessary to use the alignment of each of the studied subfamilies.

**Fig. 5.**
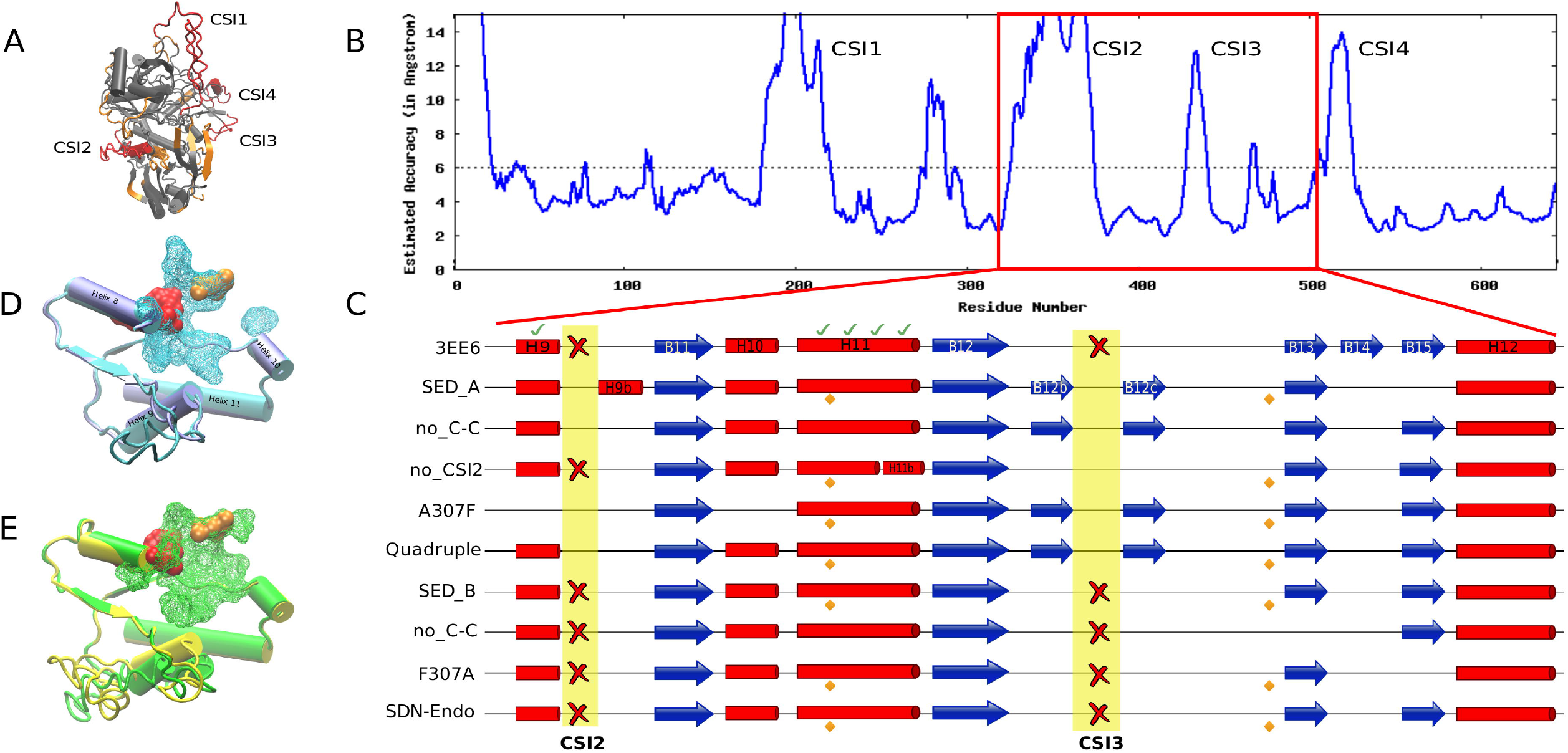
Cluster dependent Mutual Information Networks. Shown are the MI networks obtained for the hypo-endo cluster; the hypo-TPP cluster; and the combined hypo-endo / hypo-TPP cluster containing all fungal sequences. Red nodes represent the same positions in the three MI networks demonstrating high similarity between the Hypo-endo and Hypo-TPP MI networks only.

**Table 1:** Identification of SDPs

|  |  |  |  | SDPfox | MISTIC cMI |  |
| --- | --- | --- | --- | --- | --- | --- |
| Net | 3EE6 | SED_A | SED_B | Z-score | Endo | TPP |
| - | S294 | D324 | G312 | 15.5 | 235 | 412 |
| E | K346 | K407 | Q377 | 13.9 | 284 | 27.7 |
| E | V340 | Q401 | V371 | 13.8 | 244 | 173 |
| E | S73 | S76 | H75 | 12.6 | 207 | 6.5 |
| E | E343 | E404 | L374 | 9.9 | 150 | 14.1 |
| E | W307 | A343 | F336 | 8.8 | 331 | 7.2 |
| E | V89 | F92 | L92 | 8.4 | 189 | 19.1 |
| E | A349 | L410 | S380 | 8.0 | 109 | 36.7 |
| E, p | V260 | P280 | F268 | 7.2 | 114 | 25.8 |
| - | R350 | Q411 | R381 | 11.7 | 156 | 588.6 |
| - | M345 | M406 | A376 | 11.4 | 124 | 13.5 |
| - | L61 | G64 | F63 | 11.3 | 126 | 7.1 |
| T | E302 | N338 | E331 | 10.9 | 219 | 53.4 |
| T | F304 | F340 | Y333 | 7.3 | 101 | 29.4 |
| T | Q301 | L337 | N330 | 7.0 | 49 | 29.8 |
| - | L500 | V586 | M528 | 9.7 | 43 | 51.6 |
| p | Q55 | Q58 | L55 | 9.7 | 167 | 0 |
| p | A52 | G55 | L55 | 7.1 | 183 | 13.63 |
| - | L455 | I542 | Q484 | 9.5 | 123 | 21.7 |
| - | Q229 | E252 | S241 | 9.1 | 224 | 29.5 |
| - | V291 | Q321 | S309 | 8.2 | 231 | 7.1 |
| - | L80 | L83 | F82 | 7.8 | 24 | 41.5 |
| - | R506 | V592 | W534 | 7.4 | 74 | 20.9 |
| - | F51 | I54 | L51 | 7.1 | 47 | 0 |
| - | F235 | Y258 | A247 | 7.1 | 83 | 0 |

**Fig. 6.**
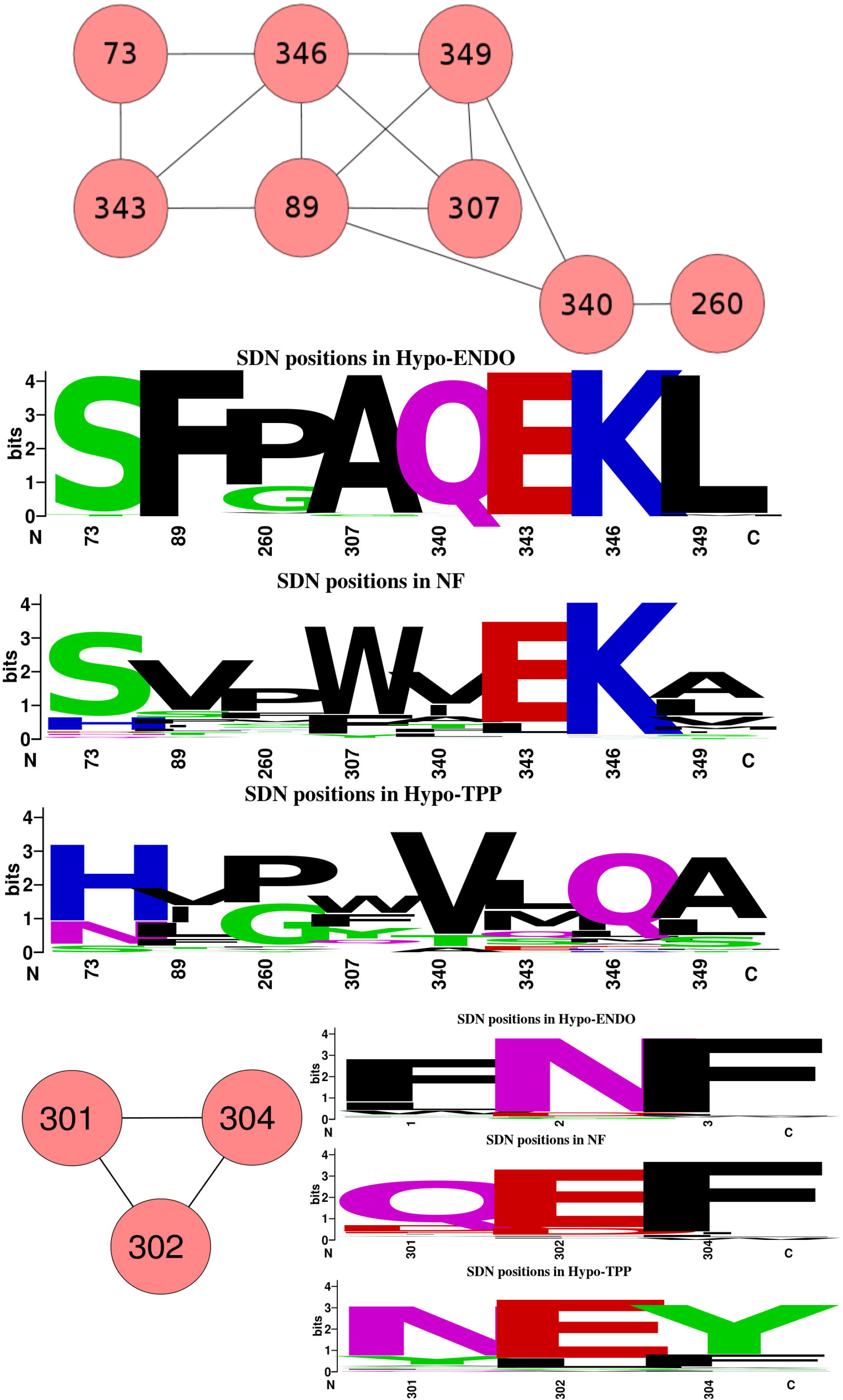
Specificity Determining Networks. SDPs, i.e. CDPs with high mutual information determined on the Hypo-Endo and Hypo-TPP subclade datasets, form one highly connected Specificity Determining Network each (numbers according to 3EE6). Sequence logos show residue conservation at the involved positions in the Hypo-Endo, the Hypo-TPP and non-fungal clades.

### 3.4 Structure-Function Prediction

Probably not all SDPs of the identified SDN will have the same impact on enzyme function. The most highly connected SDPs that are CDPs with a large physicochemical difference between the two clades, are most likely to explain the functional diversification. Both SDP346 and SDP89, located in the core and the prosegment respectively, connect to five other SDPs (Fig. 6.). Core SDP346 shows a substitution of the amphipathic, positively charged Lysine in Hypo-endo (K407 in SED_A) to most often the polar glutamine in Hypo-TPP (Gln377). In Human TPP 3EE6, K346 is located on the interface between core and prosegment interacting with E343 (Fig. 7A). Although SED_A has the same residues, K407 (homologous to K346 from 3EE6, see Table 1) physically interacts with SDP89 (F92). This, according to the model, seems to be caused by a slight dislocation of helix 9 in SED_A, on its turn forced by the additional helix 9b, which results from CSI2. The corresponding Q377 of SED_B was modeled internally and is likely stabilized in the opposite direction by the polar residue Arg 381 in SED_B (Fig. 7C) whereas F92 is replaced by a Leucine. The hydrophobic interaction between the large F92 and the aliphatic chain of K407 suggests SDP346 is involved in the interaction between prosegment and core. The ε-amino group of K407, no longer forming a salt bridge with E404 is envisaged to be protonated upon secretion into the acid environment, thereby destabilizing the interaction between core and prosegment regions. Given the suggested role of the prosegment in sedolisin folding and stabilization [16] this could explain at least part of the diversification.

**Fig. 7.**
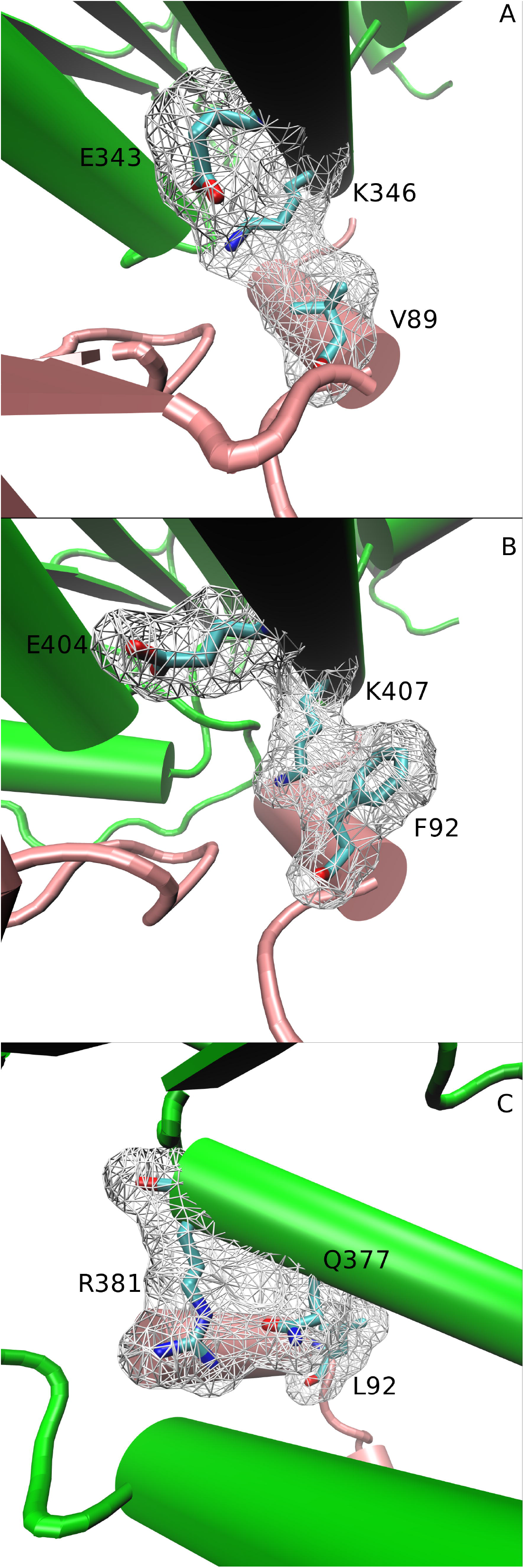
Interaction of SDP346, SDP343 and SDP89 at the interface between core and prosegment. Cartoons showing interaction in 3EE6 (A), SED_A (B) and SED_B (C). In 3EE6 K346 forms a salt bridge with E343. The different location of Helix 9 in SED_A appears to separate K407 and E404 which is compensated by F92 from the prosegment. In the SED_B model (note the different angle) Q377 is folded internally and interacts with R381. Green cartoon corresponds with core where pink cartoon represents prosegment. The SDPs are represented in licorice element, while surf represents their 5Å radius.

SDP349 also occurs in the vicinity of SDP346 and connects directly to both SDP346 and SDP89. Alanine, common in Hypo-TPP is slightly less hydrophobic than the predominant Leucine of Hypo-endosedolisins. SDP 340 is also in close range of SDP346 and connects directly to SDP89. It shows predominantly a polar Glutamine in Hypo-Endo and a hydrophobic Valine in Hypo-TPP clade. Interestingly, SDP340 is found next to position 341 that corresponds with one of two cysteines that are absent in non-fungal sedolisins and strictly conserved in fungal sedolisins (See Fig. 1.). Since strictly conserved cysteine pairs often correspond with disulfide bridges we checked their orientation in the models of SED_A and SED_B. Although in both the SED_A (S. Fig. 1D) and the SED_B model they are modeled at positions that seem to favor a disulfide bridge, this is not modeled. Nevertheless, the virtual C402A / C452A double mutant of SED_A also reverts to the structure lacking helix 9b (Fig. 4C). All together, this suggests that the SDN is related to the predicted structural changes discussed in the previous section.

SDP307, part of helix 9, is a position that, according to the MI analysis, interacts with SDPs 346, 349 and 89. Although in the structure of Human TPP 3EE6 W307 is located at over 9 Å from the local SDP network described above, in the model of SED_A, its counterpart Alanine is found at 3.7 Å (See Fig. 7B). W307 is buried in 3EE6 and present in most other non-fungal sequences and represented by an aromatic residue in the Hypo-TPP clade. Substitution of a buried aromatic residue by the small Alanine will most likely result in a conformational change. We envisaged that the substitution might be related to the conformational changes identified between 3EE6 and SED_B on the one hand and SED_A on the the other. We made structural models of virtual mutants, exchanging SED_A for SED_B residues. The model of virtual mutant A307F in SED_A suggests the loss of helices 9, 9b and 10. Compensation should, according to the above train of thought come from SDPs 89, 346 and 349, which directly connect to SDP307. Hence, we modeled the quadruple A343F-F92L-K407Q-L410S SED_A mutant. The obtained model resembles 3EE6 and SED_B since the principal helices H9 and 10 are modeled nearly identically (Fig. 4D and E).

The other SDPs show a lower level of connectivity and might involve secondary compensations. SDP 73 connects to key SDP346 as well as to SDP343. SDP 73 is mostly S in the Hypo-Endo and H/N in the Hypo-TPP clade. Structural analysis reveals that this SDP is located in the hinge region preceding the helix that contains SDP89 and following the helix that contains the pair of pSDPs 55 and 58 that we identified (S. Fig. 1.). The mentioned pair 55-58 physically interacts with SDP89 and SDP346 suggesting that they might be involved in the same diversification process. SDP73 not only might affect the interaction of SDP89 and SDP 346 but the interaction between prosegment and core in general. Finally, SDP 260 only connects to residue 340 in the SDN and is distant to the local SDP network. The fact that it concerns a proline located at a hinge position in the core, which is important in structural conformation, in Hypo-Endo sedolisins suggests this SDP might be involved in a conformational change. The Hypo-TPP network (SDP301-SDP302-SDP304) shows less pronounced physicochemical differences in residue occupation. Interestingly, it is situated within contact range of the SDP307.

## 4 Discussion

We studied eukaryotic sedolisins by computational analysis of protein sequences. The protein sequences were obtained from a large set of complete proteomes among which many of fungal origin since it has been suggested that acidification is related to expansion of the sedolisin familiy in fungi□. Therefore, the main attention of this study is directed at the evolutionary history of fungal sedolisins.

Sequences were aligned by MAFFT’s iterative refinement method and the resulting MSA was manually improved in order to correct poorly aligned hallmark residues as well as misaligned residues in secondary structure elements. Correction was likely required due to large taxonomic distances and the presence of CSIs which tend to disturb the alignment process. Prior tree reconstruction, the MSA was subjected to trimming, guided by maintaining particularly β-sheets, in order to remove these CSIs and unreliably aligned subsequences. The clustering pattern of the CSIs largely corresponds with the phylogenetic clustering (See Fig. 2B.), which confirms the topology of the tree is basically correct, as is also indicated by bootstrap support (See Fig. 2A. and B.). Interestingly, the tree contains two fungal clades and a clade with other eukaryotic sedolisin sequences. Since the fungal sequences are clearly separated from the non-fungal eukaryotic sequences it appears the evolutionary rate of fungal sedolisins has been higher than that of other sedolisins. A possible explanation for the accelerated evolutionary rate in fungi is given by the fact that fungal sedolisins are predicted to be secreted, whereas Human TPP is lysosomal. Tree topology combined with the apparent random distribution of fungal paralogs over the two fungal clades clearly suggests functional diversification, which also could result result in increased evolutionary rates. Although additional subclades can be distinguished and certain fungi have up to six paralogs, we focused on comparing the two major fungal clades based on the observation, initially made by Reichard and coworkers [23], that characterized TPPs are found in a different clade than endosedolisins (Fig. 2). We studied the role of CSIs and SDPs in the context of a conformational change observed in a model of endosedolisin SED_A, as compared to Human TPP structure and a model of TPP SED_B.

**Fig. 2.**
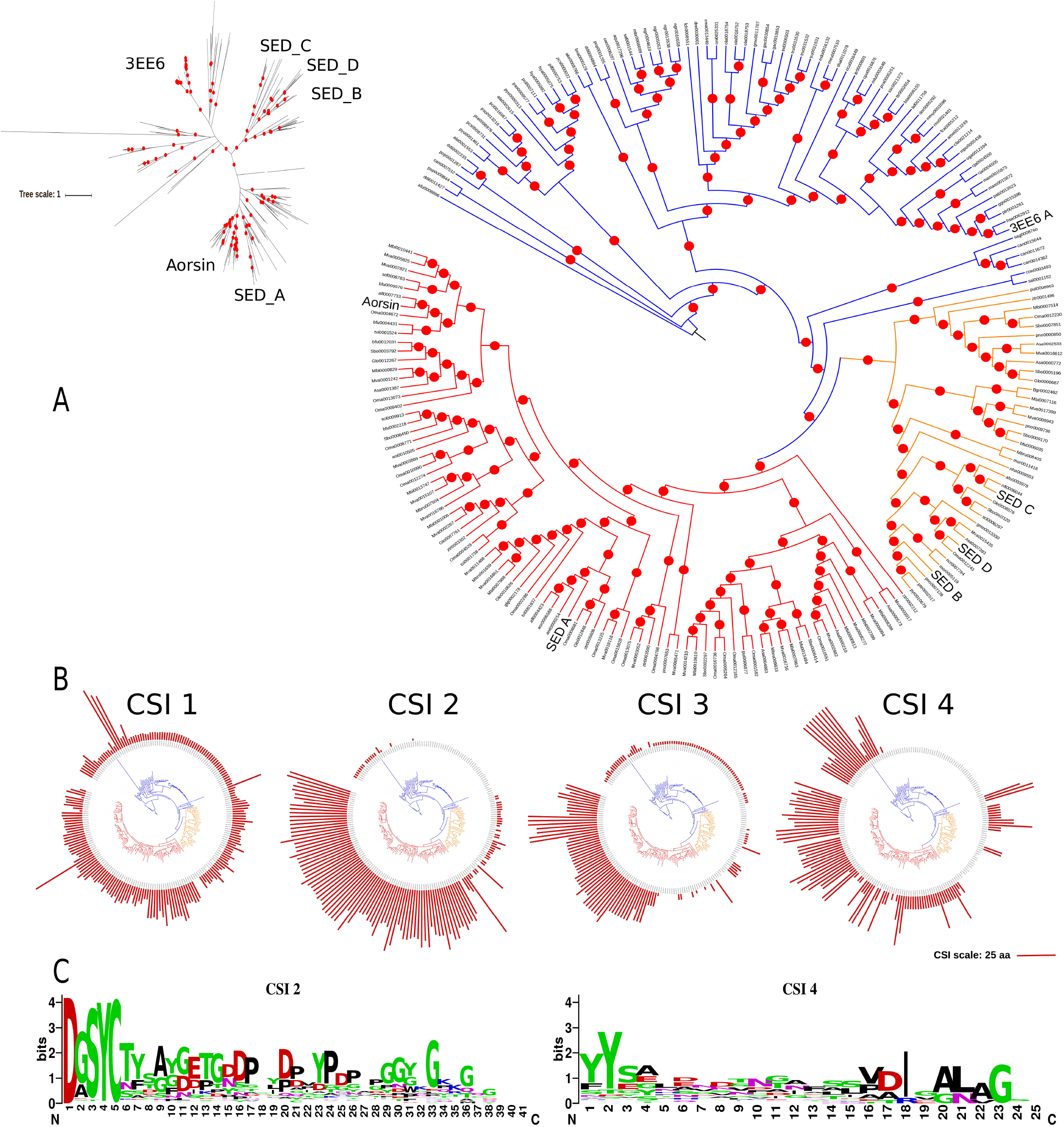
Phylogenetic Clustering and Cluster Specific Inserts of Eukaryotic sedolisins. (A) Phylogenetic trees of eukaryotic sedolisins. The large circular cladogram shows the major clades of non-fungal sequences (blue); fungal hypo-TPPs (yellow); and fungal hypo-endosedolisins (red). Red dots indicate bootstrap support of over 0.7. Sequences with known characteristics are indicated in larger font and described in the text. The small radial phylogram shows the midpoint rooted tree with distances. The scale bar indicates 1 amino acid substitution per site. (B) The circular phylograms show the clustering of the four CSIs: the length of each sequence’s CSI is represented by a red bar The scale bar at the bottom corresponds with 25 amino acid residues (C) Sequence logos of the CSI2 and CSI4 regions of the hypo-endosedolisin cluster.

The CSIs form a major obstacle when studying diversification of fungal sedolisins. Not only does their presence negatively affect the alignment process, it will also affect folding, making modeling less reliable. The residue quality (resQ) scores provided by I-TASSER are an indication of the reliability of the model and confirm that the CSI regions are unreliably modeled whereas in general confidence of the models is good (e.g. see for model of SED_A in Fig. 4A and B). The fact that the loops are likely incorrectly modeled does however not imply that their approximate location is incorrect and we envisage their solvent exposed location does not severely affect the folding of the core. It is difficult to envisage or explain how CSIs 1, 3 and 4 are related with the hypothesized diversification towards endosedolisin and TPP but the fact that CSI2 seems to instigate a conformational change and that removing CSI2 from SED_A yields a model that resembles the fold of 3EE6 (Fig. 4) is intriguing. Furthermore, CSI2 and CSI4 are conserved among all and most Hypo-endosedolisins respectively (Fig. 2B) and are located on opposite sides of the predicted pocket cavity of Hypo-endosedolisins. As such they might affect the exact conformation of the pocket. They also may have some importance in substrate binding or contain retention signals. CSI3 and CSI1 show no clear conservation and are found near to the calcium binding site and in the progsegment respectively and there are no clues regarding their functions.

A number of approaches and softwares to predict SDPs exist, although it might be noted that most often CDPs are identified. CDPs are most likely the result of positive selection but neutral evolution or even phylogenetic reconstruction artifacts can also result in CDPs. Diverge [47] should theoretically contemplate this by using likelihood models. A recent version of Evolutionary Trace [48] includes mutual information in order to substantiate its predictions. We combined SDPfox [35], to identify CDPs, with MISTIC [36], to confirm CDPs as SDPs. In a recent paper we showed that combination identified a number of known target positions in the evolution of truncated hemoglobins [49], which verifies the applicability of the method. The disadvantage of MI is the requirement of large datasets, which basically means the MI calculations in this case are not too reliable. However, since MISTIC’s z-scores tend to increase with the number of sequences and we used the default threshold, set for about 400 sequences, the MI calculation we made will most likely have suffered poor sensitivity rather than poor specificity. Then, we further extrapolated the idea that CDPs with high MI are likely SDPs towards SDP network connectivity. We identified three networks. The largest network contains seven SDPs of which two connect to five out of six other SDPs. It also contains a pSDP260. A second network identified using the endosedolisin dataset has two pSDPs. The third network contains three SDPs and is fully connected. All together, out of the 25 CDPs we selected only 10 SDPs and 3 pSDPs, corroborating the method suffers from poor sensitivity rather than poor specificity.

The advantage in using SDNs is not only reflected in that it allows for a substantiation of SDP prediction, it also relies in the fact that it shows co-evolving partners allowing a more elaborated explanation and improved predictions and understanding of the complete process of functional diversification. As stated before, based on the tree topology we envisaged a single functional diversification towards, based on biochemical evidence, endosedolisin and TPP activity. Although there are many uncertainties since it seems many factors, including SDPs but also at least one of four CSIs, are involved, all results seems to converge around the conformational change identified in the SED_A model. Models of virtual mutants directed at the central role of SDP307, the compensations of SDPs 89, 346 and 349, the predicted disulfide bridge and also the four CSIs suggest all are involved in that conformational change. Then, if we consider the conservation pattern of Trp/Phe/Tyr of SDP307 in the hypo-TPP clade, we should actually consider that position 307 is an SDP in the TPP clade. Thus, although the evolutionary processes have taken place in two clades are are therefore strictly independent, they seem to converge at a central role for SDP307. Correspondingly, SDP307 has been shown to be important for activity in Human TPP [15]. Where, in the presence of a predicted disulfide bridge C402-C452, A307 requires compensation from SDPs 340-346-349 and 89, all together resulting in a conformation that allows endosedolisin activity, W/F/Y307 would require compensation by SDPs 301-302-304 to favor TPP activity. The endosedolisin activity seems not only caused by additional helix 9b, coded by CSI2, but is also related with the interaction with the chaperone-like prosegment, as shown by the apparently important interaction between the aliphatic sidechain of K346 and F89, further strengthened by the combination of SDP73 and the pSDP pair of 55 and 58, as demonstrated in S. Fig. 1. Interestingly the pSDP pair we identified also point to the same interaction, more easily explaining the role SDP73 might play. As such we consider these two pSDPs as SDPs based on our structure-function prediction. Only pSDP260 seems difficult to explain in the same context and might well be the result of a neutral substitution given that Glycine can easily take the same conformation as a Proline.

Another consideration that can be made is how evolution has most likely occurred. Although we did not perform a prediction of the ancestral fungal sequence, a number of observations should be made. We envisage the absence of the disulfide bridge C402-C452 in the SED_A model as a a modeling artifact. The fact that two Cysteines are modeled within 5 Å distance combined with the fact that they are strictly conserved among fungal sedolisins, is not likely a mere coincidence. It can be envisaged that such a disulfide bridge stabilizes the enzyme in the oxidative extracellular environment and that the invoked structural change triggered compensations which we hypothesize have resulted in the diversification towards TPP and endosedolisin activity. This predicted bridge appears to be required for the conformational change near the binding pocket as is shown by the virtual mutant (Fig 4C). However, the presence of the disulfide bond in SED_B does not affect the conformation, suggesting its presence is not sufficient for the conformational change in SED_A. An obvious requirement is the presence of at least part of CSI2. Interestingly CSI2 is the only CSI that corresponds perfectly with the clustering into Hypo-endo and Hypo-TPP. The ancestral SDP307 was likely a Tryptophan but the other SDPs that show significant physiochemical changes seems to correspond with endosedolisins (S73, E343, K346 in 3EE6 and SED_A).

Although SDPs, mutant analysis and literature confirm each other, future validation of the proposed functional differences requires molecular dynamics and or wetlab experiments. Given the sheer amount of characteristics (i.e. CSIs and SDPs) involved, this will be difficult to achieve. As such, maybe a more feasible approach is to obtain a structure of SED_A or another characterized endosedolisin such as Aorsin.

The remaining question concerns how this change in conformation is related to the difference in activity. TPP activity can be envisaged to require a less spacious binding cleft, as can be seen by comparing Fig. 4D and E, in concordance with the fact that SDP307 is occupied by a large hydrophobic residue in the Hypo-TPP and NF clades and by small hydrophobic in Hypo-Endo.

## Legends Supplemental Figure

**S. Fig. 1.** A, B and C (3EE6, SED_A and SED_B respectively: Interaction between pSDPs 55 and 58 and the major SDN via SDP73, afecting the interaction between prosegment and catalytic domain. D Modeling of Cysteines 402 and 452 from SED_A, strictly conserved among all fungal sedolisins and absent in non-fungal sedolisins.

